# In situ structural analysis of SARS-CoV-2 spike reveals flexibility mediated by three hinges

**DOI:** 10.1101/2020.06.26.173476

**Authors:** Beata Turoňová, Mateusz Sikora, Christoph Schürmann, Wim J. H. Hagen, Sonja Welsch, Florian E. C. Blanc, Sören von Bülow, Michael Gecht, Katrin Bagola, Cindy Hörner, Ger van Zandbergen, Shyamal Mosalaganti, Andre Schwarz, Roberto Covino, Michael D. Mühlebach, Gerhard Hummer, Jacomine Krijnse Locker, Martin Beck

## Abstract

The spike (S) protein of severe acute respiratory syndrome coronavirus 2 (SARS-CoV-2) is required for cell entry and is the major focus for vaccine development. We combine cryo electron tomography, subtomogram averaging and molecular dynamics simulations to structurally analyze S *in situ*. Compared to recombinant S, the viral S is more heavily glycosylated and occurs predominantly in a closed pre-fusion conformation. We show that the stalk domain of S contains three hinges that give the globular domain unexpected orientational freedom. We propose that the hinges allow S to scan the host cell surface, shielded from antibodies by an extensive glycan coat. The structure of native S contributes to our understanding of SARS-CoV-2 infection and the development of safe vaccines. The large scale tomography data set of SARS-CoV-2 used for this study is therefore sufficient to resolve structural features to below 5 Ångstrom, and is publicly available at EMPIAR-10453.

## Introduction

As of June 2020, the severe acute respiratory syndrome coronavirus 2 (SARS-CoV-2) pandemic has reached all five continents with over >9 million confirmed cases and >460.000 SARS-related deaths. Compared to its close relatives SARS-CoV and MERS-CoV, SARS-CoV-2 appears to spread more efficiently. Without a vaccine, human distancing is currently the only effective measure to contain the pandemic, causing dramatic impact on the socio-economical landscape. The coronaviral spike surface protein (S) is the primary vaccine target, being exposed on the virus, mediating binding of the virus to the cell surface and fusion with cellular membranes required to initiate infection (*de Haan and Rottier, 2005*). S also determines tissue- and cell tropism and mutations in S may alter the host range of the virus and cross species barriers *(Hulswit et al., 2016; Li, 2016*). Vaccine efforts therefore focus on inducing cellular and humoral immunity with neutralizing antibodies that block viral binding, entry, fusion and infection by binding to S.

S is a trimer that adopts a typical club-like shape of ∼20 nm in length on the surface of the virus. The ectodomain consists of a globular top and a slender stalk that connects the former to the membrane. Because S is essential for infection and the major target of vaccine development, understanding its molecular structure is of utmost importance and truncated or detergent extracted S over-expressed in mammalian cells has been subjected to various structural investigations *in vitro* by single particle cryo-EM (Wrapp et al., 2020, Walls et al., 2020, Cai et al., bioRxiv 2020, Henderson et al., bioRxiv 2020, Xiong et al., bioRxiv 2020). SARS-CoV-2 S is structurally similar to SARS-CoV S and binds to the same receptor, the angiotensin-converting enzyme 2 (ACE2) (Walls et al., 2020, Wrapp et al., 2020). The globular domain contains the receptor binding domains (RBDs) that reside on top and are shielded by the N-terminal domains (NTDs). Two conformational states have been observed for the truncated versions of S investigated *in vitro*. A closed conformation in which all three RBDs are positioned in-between the NTD and the RBD of the neighbouring monomer, thereby hiding the receptor binding site; and an open conformation in which one of the RBDs is exposed upwards, while the other two remain in the closed conformation (Wrapp et al., 2020, Walls et al., 2020). Although it has been suggested that this conformational variability might be related to receptor binding (Wrapp et al., Science 2020), *in vitro* structural analysis of detergent extracted, full-length S exclusively observed the fully closed conformation (Cai et al. bioRxiv 2020). The latter study also reported a spontaneous transition into the post-fusion state during purification. The extent to which such spontaneous transitions occur on the viral surface remains uncertain. Further, the most C-terminal part of the protein, in particular large parts of the stalk and the entire transmembrane domain remained poorly resolved in any of the previous studies.

In some coronavirus strains, S may be cleaved at the interface between the S1 and S2 subunits by furin, a cellular protease located in late or post-Golgi compartments; the N-terminal S1 harbors the RBD, while the C-terminal S2 contains the fusion machinery and the transmembrane anchor. Although not necessary for infection, furin cleavage of S is linked to virulence *in vivo (Li 2016; Belouzard et al., 2012)*. Considerable effort has been undertaken to stabilize S for structural investigations *in vitro (Walls et al., 2017b)*. To lock S in a pre-fusion conformation the furin cleavage site was deleted and two proline residues introduced at position 986 and 987 *(Kirchdorfer et al. 2018)*. Further, the truncated ectodomain of S was fused to an artificial C-terminal trimerization tag *(Wrapp et al., Science 2020; Walls et al., 2020*; *Xiong et al*., bioRxiv *2020*). To potentially facilitate the generation of conformation specific antibodies, dedicated constructs of S have been engineered that lock the RBDs into either the open or closed conformation *(Henderson et al., bioRxiv 2020, Xiong etal., bioRxiv 2020)*. To what extent these conformational states are representative of the *in situ* scenario on the viral surface remains unknown.

The SARS CoV2 S is a type I membrane glycoprotein with an N-terminal signal sequence for synthesis on, and translocation across the endoplasmic reticulum (ER), where it trimerizes and becomes glycosylated at its 22 predicted N-linked glycosylation sites. S protein transport through the Golgi apparatus and further processing of its glycosylation is thought to occur once incorporated into virions that bud at pre-Golgi membranes and exit infected cells by vesicular transport (*de Haan and Rottier, 2005*). Previous *in vitro* cryoEM studies resolved roughly two thirds of the predicted 22 N-linked glycans (Walls et al., 2020; Wrapp et al., 2020). Although an N-terminal signal peptide for secretion was fused to S, it is not clear if over-expression of the protein alone in HEK 293F cells (Walls et al., 2020) ultimately led to the same glycosylation pattern as compared to viral assembly that massively impacts onto various organellular systems (Gordon et al., 2020; Bojkova et al., 2020; Ulasli et al., 2010).

While these previous studies provided valuable data of isolated spike protein or its fragments, *in situ* structural analysis of SARS-CoV-2 remains preliminary. Structural information about S *in situ* is a prerequisite towards understanding the molecular mechanisms of host cell entry. It would allow to explore the conformational variability within the virion and the accessibility of epitopes for neutralizing antibodies; to determine the glycosylation pattern of S generated during viral assembly; and to analyze the yet unresolved stalk domain that flexibly connects S to the viral surface. Here we combined cryo electron tomography of purified SARS-CoV-2 virions with subtomogram averaging, molecular dynamics simulations and integrative structural modeling to address these challenges.

## Results

### Large scale tomographic analysis of SARS-CoV-2 virions

To structurally analyze SARS-CoV-2 spike protein *in situ*, we passaged the virus through tissue culture cells and purified it from the inactivated supernatant by sucrose centrifugation (see methods for detail). The origin of the SARS-CoV-2 isolate used in this study is south Germany and described in detail in the supplement. We acquired a large scale cryo electron tomography data set of purified virions using acquisition parameters that are suitable for subtomogram averaging (see methods). Visual inspection of the tomographic reconstructions revealed a very high quality data set in which, individual protein domains are clearly visible (Figure 1A, Movie S1). The raw data of 266 tilt-series with >1000 viruses have been deposited EMPIAR-10453 and might be used in the future to explore various structural properties of SARS-CoV-2, in particular its pleiomorphic features. In this initial study we will exemplify the potential of our data as a resource with the *in situ* structural analysis of S.

**Figure 1.**
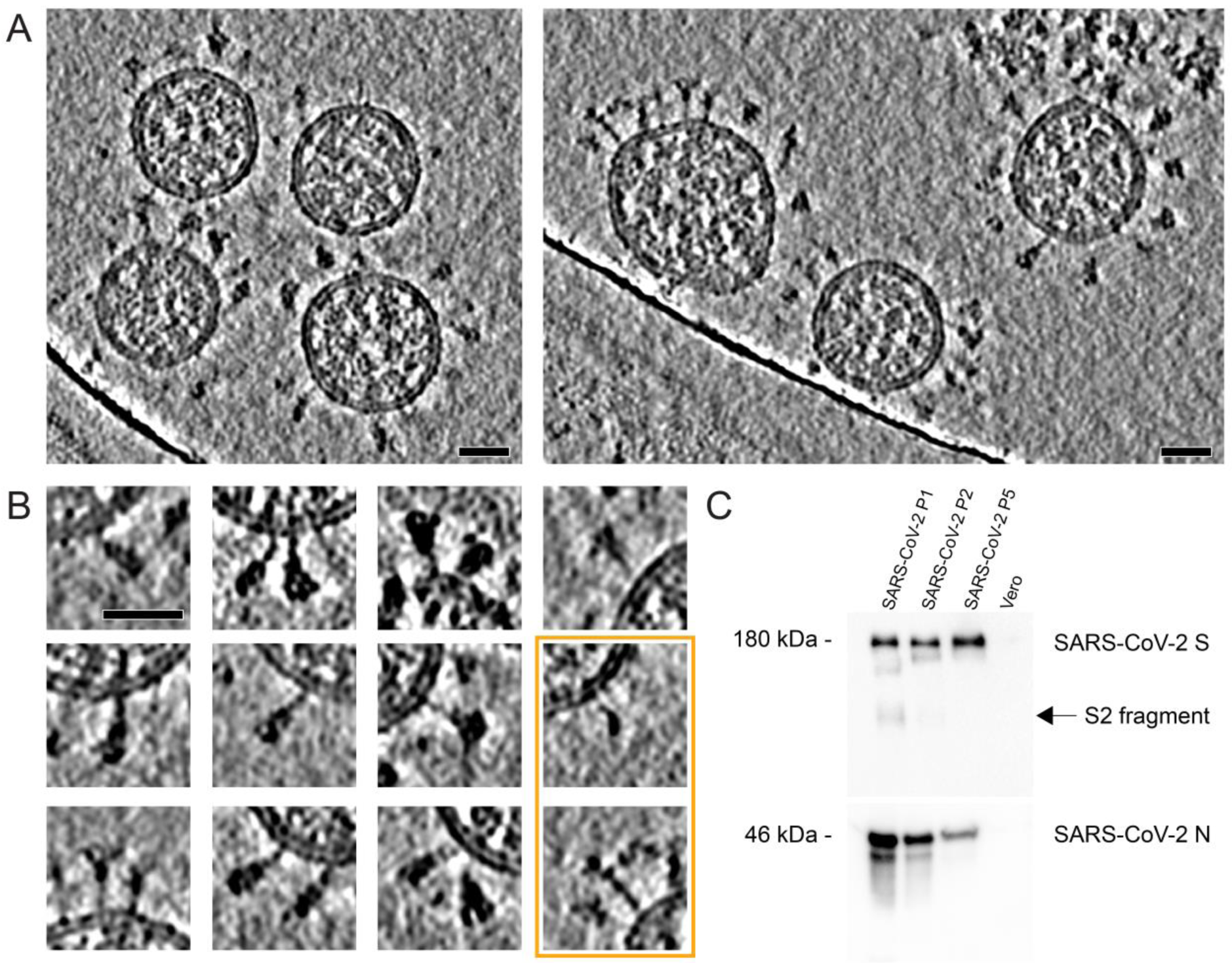
Cryo electron tomography of SARS-CoV-2 virions. **(A)** Slices through tomographic reconstructions of highly pleiomorphic SARS-CoV-2 virions. Scale bar 30 nm. **(B)** Same as (A), but as a gallery highlighting individual copies of S. All domains, including the transmembrane part, are clearly resolved. Although the vast majority of S is reminiscent of the pre-fusion conformation, the particles framed in orange are strikingly similar to the post-fusion conformation as proposed by (Cai et al, bioRxiv 2020; compare to their Figure 4C). **(C)** Western blot showing the loss of furin cleavage products of SARS-CoV-2 S and the increase of uncleaved S (180 kDa) within 5 passages through tissue culture (loading control using anti-N antibody).

### The pre-fusion conformation of spike dominates the surface of virions purified from tissue culture cells

Visual inspection of viral surface revealed that spike was mostly present in the pre-fusion conformation, while some notable exceptions appeared strikingly similar to previous analyses of an alternative conformation (Figure 1B) thought to be the post-fusion conformation (Cai et al bioRxiv 2020; Duquerroy *et al*., 2005). We hypothesized that the pre-fusion conformation might have been positively selected during passage through tissue culture cells, in the virions that we purified from the supernatant. We sequenced the viral genome at passages 1 to 5, the latter of which was used for the preparation of cryo-EM samples. The furin proteolytic cleavage site was gradually lost during passage through tissue culture as shown by Sanger sequencing and western blot analysis on the protein level confirming previous studies (Figure 1C, Figure S1; Lau et al., bioRxiv 2020; Ogando et al., bioRxiv 2020). It has been previously noted that due to the location of the furin site within a loop, its cleavage may have little impact on the S trimer in the pre-fusion conformation, other than ultimately allowing for the dissociation of the S1 fragment (Cai et al., bioRxiv 2020). A study that appeared on bioRxiv during the preparation of this manuscript suggests that in virions produced in Vero E6 cells the pre-fusion is predominant while this may be different for other cell types (Klein et al., bioRxiv 2020), thus underscoring that our large scale data set is representative of the native state.

We manually picked the virions and oversampled their exterior to extract subtomograms (see methods for detail). After a few iterations of alignment, they converged into a low-resolution structure of the globular domain of spike in its pre-fusion conformation. Statistical analysis of the resulting positions revealed that on average 40 copies of S trimer reside on the surface, although their density varies across virions. Individual spike complexes appeared randomly distributed on the viral surface and did not show any significant tendency to cluster but rather had slightly repellent properties (Figure 2A).

**Figure 2.**
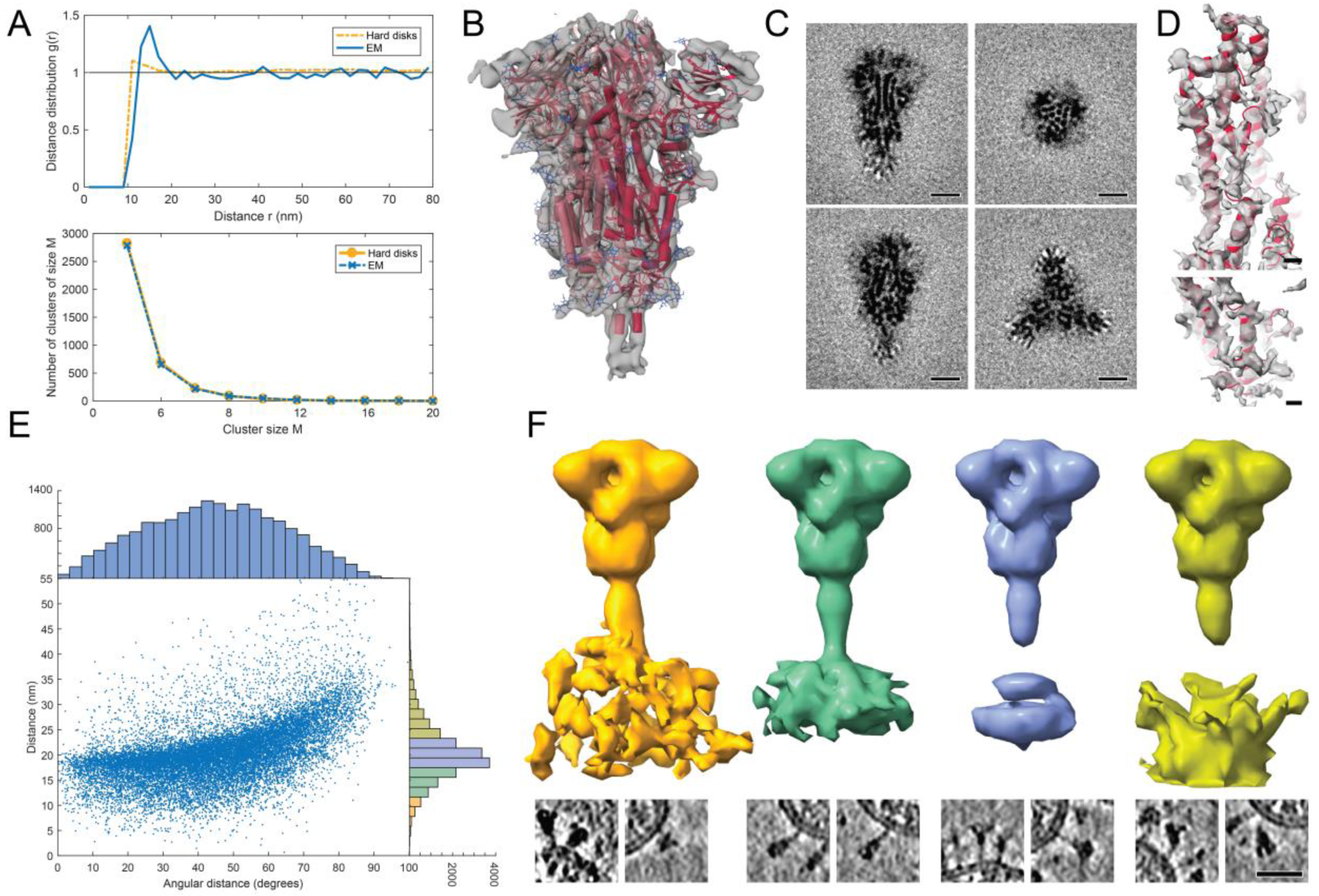
Subtomogram analysis of SARS-CoV-2 S protein. **(A)** Distance (top) and cluster-size distributions (bottom) of S on the viral surface, with non-overlapping hard disks of 10 nm diameter as reference. **(B)** Subtomogram average of the ectodomain of S shown isosurface rendered and fitted with the previously published atomic model as determined by single particle EM (PDB ID 6VXX). Subtomogram average in transparent grey, secondary structure elements in red and glycosylation sites in blue. **(C)** Same subtomogram average but shown as slices through the reconstruction. Scale bar 5 nm. **(D)** Detail of the average of the symmetric unit of S. Scale bar 5 Å. **(E)** Distribution of the angular orientation and distance of the ectodomain with respect to the bilayer. **(F)** Averages of subtomograms classified according to the distance of the ectodomain (increasing from left to right) from the bilayer. Examples of individual particles are shown as slices at the bottom panel (scale bar 30 nm). At an optimal distance, the stalk domain stretches out and is resolved.

We refined the globular domain to 7.6 Ångstrom using procedures implemented in NovaSTA (https://github.com/turonova/novaSTA) and STOPGAP packages (https://github.com/williamnwan/STOPGAP, see methods for detail). At this resolution, secondary structure elements and individual glycosylation sites are clearly discernible (Figure 2B, C). Although there was no symmetry imposed during the refinement, the average was highly symmetric, suggesting that conformational variability across the three different monomers is minimal. Structural analysis of the asymmetric unit yielded an average map from 65780 subtomograms at an overall 4.9 Ångstrom resolution. In particular the cluster of parallel helices in the center of the globular domain was nicely resolved (Figure 2D) and flexible fitting revealed that larger side chains are clearly discernible (Figure S3). Simulated annealing-based classification as implemented in STOPGAP of both, the S-trimer but also the asymmetric unit did not yield any distinct classes. However, separate cross-correlation against the single particle structures with either the closed or open RBD identified a fraction of S-trimers in the open conformation (Figure S4).

### An unusual coiled-coil resides next to a hinge in the stalk domain of spike

While the globular domain was fully contained in the tomographic map, only the directly adjacent part of the stalk domain was resolved, emphasizing the flexibility of the latter. Remarkably, a right-handed coiled coil emerges from the neck of the spike head (∼P1140-L1159), forming the upper part of the stalk domain (subsequently referred to as the “upper leg” of the stalk). Right-handed trimeric coiled coils were long thought to be absent from the structural proteome (Harbury *et al*., 1998), but can be seen in the post-fusion structure of the related mouse hepatitis virus (MHV) spike (PDB ID 6B30; Walls *et al., 2017a*). The density in the single-particle structures of SARS-CoV-2 spike (Wrapp *et al*., 2020), albeit only moderately resolved in this region next to the trimerization tag, is consistent with a right-handed coiled-coil. The assignment as right-handed coiled-coil is also consistent with the observation that this sequence stretch is not part of the post-fusion left-handed coiled-coil, as resolved for SARS-CoV spike (PDB ID 1WYY; Duquerroy *et al*., 2005), which indicates a certain incompatibility with left-handedness.

The observed electron density sharply declined at a defined position within the trimeric coiled-coil, suggesting at least one flexible hinge (Figure 2B). We analyzed the conformational variability of the stalk domains based on the position of the globular domain and its orientation with respect to the membrane as defined by subtomogram averaging. The globular domain exhibited large positional and orientational freedom, tilting up to ∼90 degrees with respect to the normal at distances of 5-35 nm from the membrane without discernible correlation (Figure 2E). We grouped our subtomograms into four classes according to their distance from the bilayer and averaged them separately. In most classes the stalk remained invisible and the density of the bilayer was blurred, suggesting a highly kinked conformation of the stalk (Figure 2F). At an intermediate distance, however, parts of the stalk and bilayer were resolved, suggesting a more defined conformation. We sub-selected ∼3200 particles in which the globular domain was oriented roughly perpendicular to the membrane and averaged them separately (see methods). In the resulting average, the stalk domain was resolved (Figure S5A). Visual inspection of the respective subtomograms, in which the stalk domains are clearly observed, further corroborated the idea of a kinked stalk with potentially several hinges (Figure 2F). We used local masking to focus the subtomogram averaging on the lower part of the stalk domain (subsequently referred to as the lower leg). We obtained a moderately resolved structure that would be consistent with the continuation of the coiled coil below a flexible hinge (subsequently referred to as the “knee”, Figure S5B).

### Molecular dynamics simulations of spike in the context of the bilayer

We performed a 2.8 microseconds long all-atom molecular dynamics (MD) simulations of a 4.1-million atom system containing four glycosylated S proteins anchored into a patch of viral membrane and embedded in aqueous solvent (Figure 3A). The simulations helped us pinpoint the molecular origins of the flexibility seen in the tomograms. Whereas the S head remained stable, the stalk exhibited pronounced hinging motions at the junctions between S head and upper leg (“hip”), between upper and lower leg (“knee”) and between lower leg and transmembrane domain (“ankle”), consistent with pronounced leg segments seen in raw tomograms (Figure 3B-C). The hip joint flexed the least (16.5 +/-8.8 deg), followed by the ankle (23.0 +/- 11.7 deg) and knee (28.4 +/- 10.2 deg) (Figure 3D).

**Figure 3.**
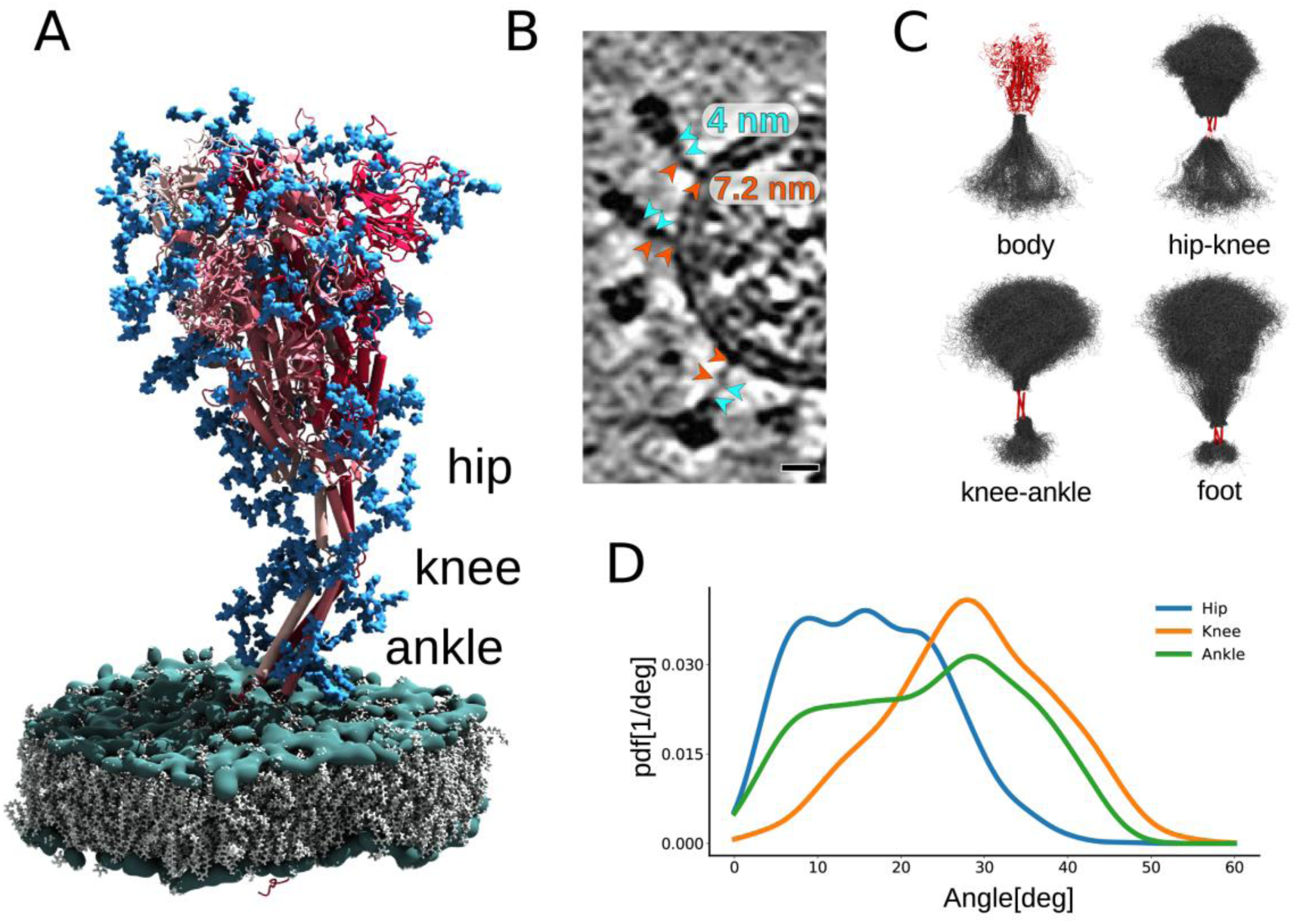
MD simulations of SARS-CoV-2 S protein. **(A)** Model of the S protein. The three individual chains of S are shown in shades of red, N-glycosylation in blue, lipids of the ER-like membrane in grey with phosphates in green; “hip”, “knee” and “ankle” mark positions of the three flexible hinges. **(B)** Examples of the hinges as seen in the de-convoluted tomograms. Cyan and orange arrowheads indicate upper and lower leg respectively, with their typical lengths indicated. Scale bar 10 nm. **(C)** Hinge flexibility in the MD simulation illustrated through backbone traces (grey) at 75 ns intervals with different parts of the S protein fixed (red). **(D)** Probability density functions for hinge bending angles at hip, knee and ankle.

### Hinges predicted by MD simulations are consistent with the tomographic data

The 3D tomographic density of S proteins protruding from the viral surface is well described by selected MD snapshots even without flexible fitting (Figure 4A). Strikingly, hip, knee and ankle can be pinpointed in individual copies of S seen in the tomographic reconstructions (Figure 4A). We flexibly fitted suitable snapshots of the MD simulations into subtomogram averages classified according to the distance of the globular domain from the membrane (compare Figure 2F to Figure 4B). In the stalk region, hinge bending gives S the required flexibility. The hinges facilitate a lateral displacement of the upper and lower leg with respect to the globular domain, if the latter is positioned closer to the membrane. As a result the stalk is diluted out in subtomogram averages focused on the globular domain (Figure 2B-C, F, 4B) but can be retrieved if the globular domains are aligned with the membrane normal (Figure S5A) or if the stalk itself is averaged separately (Figure S5B). We calculated electron density averaged over the whole MD trajectory and filtered it to a resolution comparable to the classified subtomograms. We obtained a highly similar 3D map that underscores this interpretation (Figure 4B). In rare cases, the coiled-coil near the membrane appears to be unfolded in the original tomograms (Figure S5C) and continuous with the disordered loops of the MD model.

**Figure 4.**
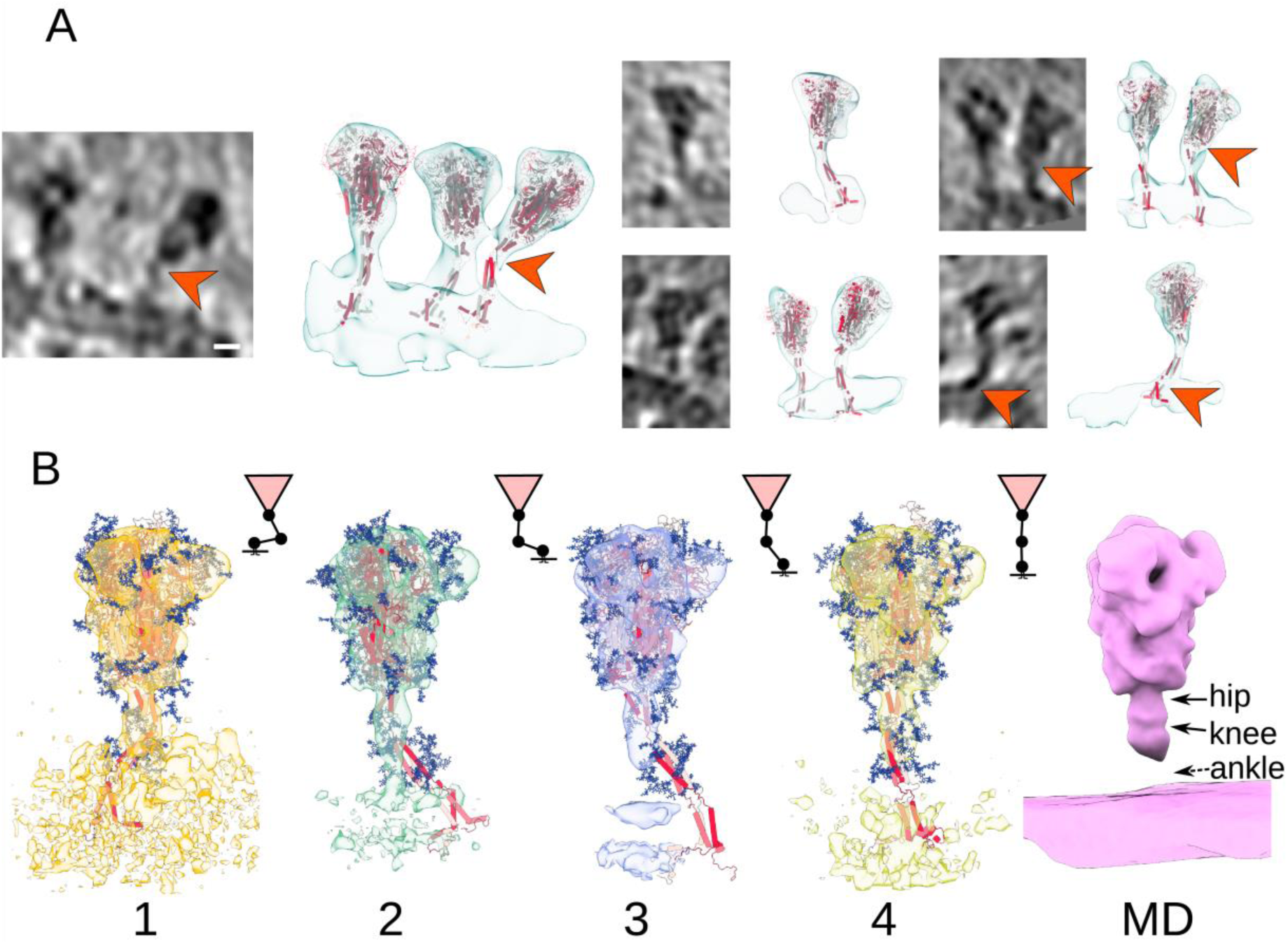
Fitting of molecular simulations into cryo electron tomograms. **(A)** The hinges of the stalk domain predicted by structural modeling (orange arrowheads) are consistent with the tomographic data. Slices through tomograms (left) and snapshots of respective MD simulations superimposed with isosurface rendered tomograms are shown for multiple instances. Scale bar 5 nm. **(B)** Fit of snapshots of MD simulations into the classes obtained for different distances of the globular domain from the membrane (1-4) as presented in Figure 2F. Shorter distances are concomitant with a stronger bending of the hinges and a lateral displacement of the stalk. Average MD density filtered to a resolution comparable to the subtomogram averages is shown as isosurface render (right).

### N-glycosylation sites are very pronounced in tomographic data

The predicted N-glycosylation sites, many already annotated in single particle EM maps (Walls et al., 2020), are generally very pronounced in the subtomogram averages. Aided by the MD model, we systematically examined glycosylation of S. The electron density of N-glycans averaged over the MD trajectory was highly consistent with the tomographic map (Figure 5A). Clustered glycosylation sites are visible in the raw electron density before averaging, e.g. protruding from the lower part of the S body (Figure 5B). Analysis of individual sites in subtomogram averages further supports the decoration of spikes with rather large glycan chains (Figure 5C). Notably, a number of sequons were resolved with more pronounced branching than previously reported in the single particle maps (Walls et al., 2016). By contrast, the density map in the region of two previously reported O-glycosylation sites (Shajahan et al., 2020) does not allow for unambiguous assignment (Figure S6A), suggestive of low glycan occupancy at these sites *in situ*. Sequon N17LT, due to its location on the unstructured N-terminus, was not localised in the density (Figure S6B). Elongated features protruding from the tip of the N-terminal domain (Figure S6B) may suggest presence of sequons N74GT and N149KS, but are less pronounced as compared to the N-glycosylation sites discussed above, likely because they are located on highly flexible loops. All these sites were recently shown to be in a close proximity to an antibody bound to the N-terminal domain (Chi et al., 2020).

**Figure 5.**
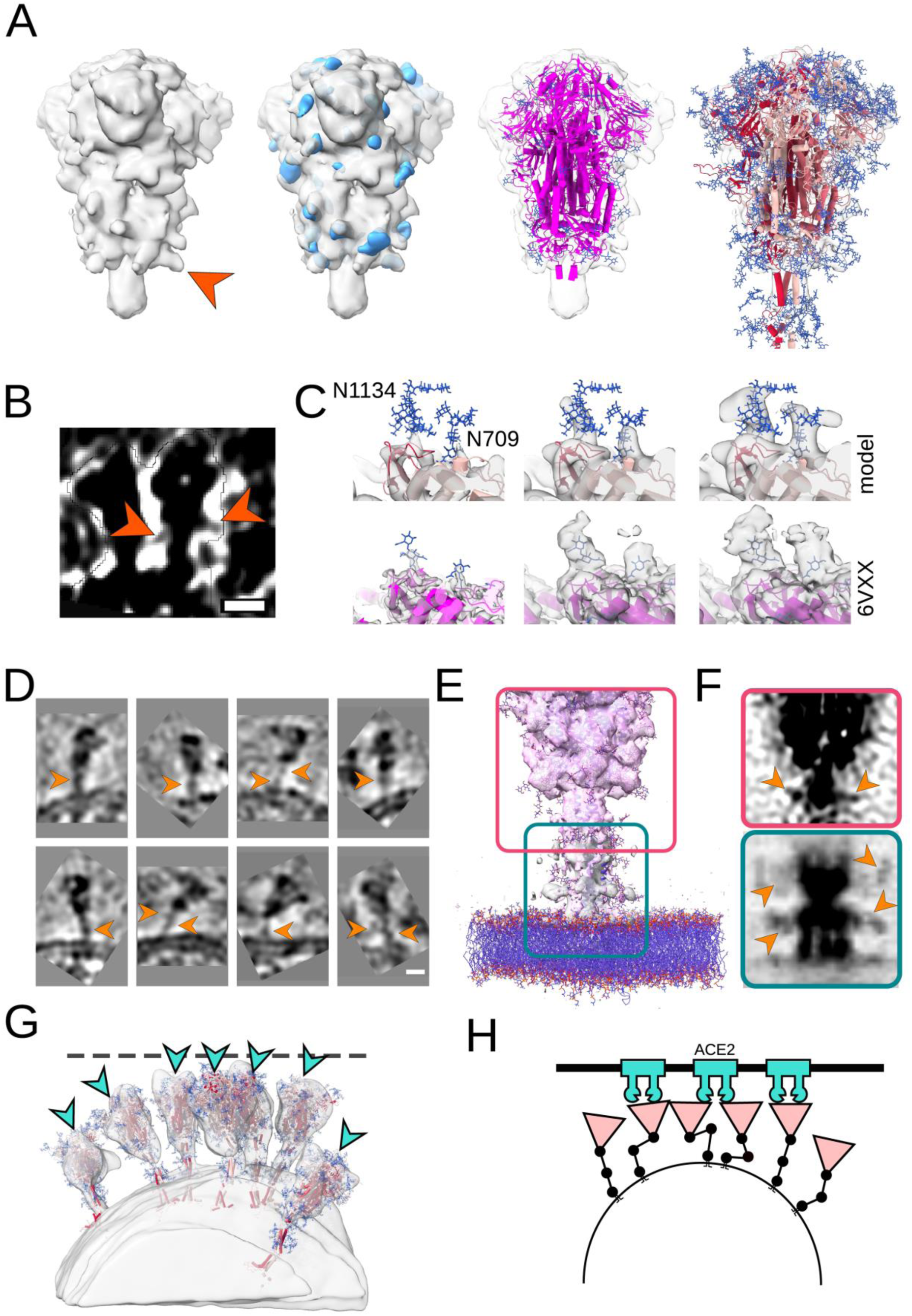
Analysis of S protein glycosylation sites and epitopes. **(A)** N-glycosylation sites are clearly discernible in the subtomogram average of the globular domain. From left to right: Isosurface rendering of subtomogram average with an individual N-glycosylation site indicated (orange arrowhead); same superimposed with the MD-calculated density for all annotated N-glycosylation sites; superimposed with previous structural model of the globular domain (PDB ID 6VXX); superimposed with a snapshot of the MD simulations. N-glycosylation sites are shown in blue. **(B)** Tomographic slice highlighting an N-glycosylation site (orange arrow heads) in the original data. Scale bar 5 nm. **(C)** Highlight of N-glycosylation positions 709 and 1134 of the MD simulations (top) and in a previous structural model (bottom; PDB ID 6VXX, EMD-21452). The subtomogram average is shown superimposed at different isosurface thresholds (transparent grey). Extensive additional density is visible. **(D-F)** The stalk domain is heavily glycosylated at the hinges. **(D)** Exemplifying tomographic slices with bulky density at the hinge positions (orange arrowheads). Scale bar 5 nm. **(E)** Superposition of the subtomogram averages (transparent grey isosurfaces) of the globular domain (framed red) and the stalk domain (framed green) with a respective snapshot of the MD simulations emphasizing the glycosylation at the hinges. **(F)** Same as (E) but shown as maximum intensity projection through the subtomogram averages. **(G)** Fits of snapshots from MD simulations into the surface of a virion; tomogram shown isosurface rendered in transparent grey. The position of epitopes for neutralizing antibodies at the RBDs are indicated with cyan arrowheads. **(H)** Cartoon illustrating a hypothetical docking event in which the hinges facilitate the engagement of multiple instances of S with their receptors.

### Integrative analysis suggests that the hinges are heavily N-glycosylated

N-glycosylation is predicted also on the knee (N1158HT, N1173AS) and the ankle (N1194ES) outside the region resolved by single particle EM (Figure 3A). We observed that these positions generally appear more bulky in tomographic reconstructions than one might expect if they were not glycosylated (Figures 1B and 5D). In subtomogram averages of the globular domain and stalk, additional density is very clearly observed at the respective positions (Figure 5E, F and Figure S5A, B), strongly suggesting that they are modified. Consistent features are present in the electron density calculated from the MD trajectory aligned on the lower leg (Figure S6C), where we can unequivocally attribute them to glycans. N-glycosylation in this position might protect the functionally important hinges from antibody binding and help to keep them flexible.

## Discussion

The two major structural analysis techniques combined in this study are complementary. While cryo electron tomography facilitates the experimental analysis of cryo fixed specimen at moderate resolution in their native context, molecular dynamics simulations reveal protein motions at atomic resolution as a function of time. In the particular case of the SARS-CoV-2 spike protein, the tomographic data reveal its conformational freedom and glycosylation sites on the viral surface but they cannot resolve the flexible stalk domain to high detail. The molecular dynamics simulations integrate various experimental parameters, in particular single-particle structures and independently determined glycosylation sites, and allow fully atomic models to evolve in time.

Our MD simulations revealed three highly flexible hinges within the stalk, coined hip, knee and ankle, which are consistent with the experimental data. One might speculate that the high degree of conformational freedom of the spike on the viral surface is important for the mechanical robustness of the virus. It might also allow the spike to engage the relatively flat surface of host cells with better avidity prior to fusion or find access to its receptor more efficiently in the crowded environment of tissue *in vivo* (Figure 5G-H). This would allow the virus to explore the surface of host cells more effectively and potentially to bind to multiple copies of ACE2. Tomographic studies of actual infection events might further address this in the future.

In contrast to the pre-fusion conformation of spike, the post-fusion conformation previously observed *in vitro* and *in situ* (Cai et al., bioRxiv 2020 Klein et al., bioRxiv 2020) and in this study (Figure 1B) is entirely straight and apparently inflexible. A particularly unusual feature masked at the edge of the resolved density of the single-particle structures but well resolved in the subtomogram averages is the short right-handed coiled-coil at the top of the pre-fusion stalk. Being lost in the post-fusion structure as resolved for SARS-CoV (PDB ID 1WYY; Duquerroy et al. 2005), we speculate that the right-handed coiled-coil is only marginally stable, priming the protein for a large structural reorganization in a spring-loaded viral fusion mechanism. These findings underline that all three hinges become irrelevant during the transition to the post-fusion conformation, which places them outside the structural core (Cai et al., bioRxiv 2020; Duquerroy et al., 2005).

A remarkable difference between SARS-CoV and SARS-CoV-2 spike is the presence of a furin cleavage site in the latter (Wrapp et al., 2020). This site is rapidly lost by passaging in cell culture as shown before (Lau et al., bioRxiv 2020; Ogando et al., bioRxiv 2020) likely because its loss favours virions with stable spikes. In Vero cells this results in the release of SARS-CoV-2 virions with S in a closed formation (this study; Klein et al., bioRxiv 2020). In the latter recent study the authors proposed that S released on SARS-CoV-2 particles is readily transformed into a conformation resembling the post-fusion protein when exposed to cells that express high levels of ACE2. Whether this conformational change relates to receptor binding, to cleavage at the S2’ site by the serine protease TMPRSS2 (Belouzard et al., 2012) or to both was not addressed in this study. In addition, it was not clear if the furin cleavage site is intact in the unpassaged isolate used in this study and, if so, to what extent S was cleaved during passage of newly assembled virus through the secretory pathway. In Vero cells we found for instance cleavage of S containing the furin cleavage site to be partial, with the majority of S remained unprocessed, suggesting that efficiency of furin cleavage may be cell type dependent. Nonetheless, the observations made in this study are interesting and open the possibility to compare the pre-fusion with the presumed post-fusion structure of S *in situ*.

The predominant *in situ* conformation of S observed in our study is the closed pre-fusion conformation. This finding emphasizes that the highly-engineered, recombinant versions of S in which this conformation is locked (Henderson et al., bioRxiv 2020, Xiong bioRxiv 2020) may indeed be valuable tools for vaccine development, although there are also differences to the *in situ* structure. N-glycosylation sites appear very bulky in the tomographic map as compared to previous single particle analysis, suggesting that decoration with sugars may indeed be more extensive during viral assembly as compared to the recombinant ectodomain modified during default vesicular transport. Our map is suggestive of additional N-glycosylation at the hinges of the stalk domain and possibly on the very tips of S NTDs. This might be important for the accessibility of epitopes on the crowded viral surface were the NTD and stalk domains appear occluded by neighbouring spikes (Figure 5G). Indeed, glycan shielding of epitopes on the coronavirus spike protein has been described before (Walls et al., 2016) emphasizing the importance of identifying the glycosylation pattern on the native spike protein. We did not detect any density for the predicted O-glycosylation sites emphasizing that it must be less extensive as compared to N-glycosylation.

Here we demonstrate the potential of our large scale tomographic data set for resolving structural features of SARS-CoV-2 particles to high resolution in their native context. The *in situ* structure of several key viral components, including e.g. the M protein that is highly enriched in the membrane and the nucleocapsid remain enigmatic. Our data might thus be explored to resolve the structures of such features in the future. We further demonstrate that high resolution structural models can be fitted directly into the tomographic reconstructions, underlining the remarkable quality of the data. This strategy might be further explored in the future to build structural models of entire virions.

## Depositions

The original tilt series have been deposited (EMPIAR-10453). Subtomogram averages were deposited into the electron microscopy data base under accession numbers EMD-11222 (S-trimer), EMD-11223 (asymmetric unit with closed RBD), and EMD-11347 (asymmetric unit with open RBD).

## Author contributions

B.T. experimental design, tomographic reconstruction, particle picking, subtomogram averaging, structural analysis, paper writing. M.S. molecular dynamics simulations, structural analysis, paper writing. C.S. experimental design, virus purification and biochemical analysis. W.H. experimental design, cryo-EM data acquisition, tomographic reconstruction. S.W. experimental design, sample preparation and screening, data analysis. F.E.C.B., S.v.B., M.G. and R.C. molecular dynamics simulations, structural analysis. K.B. experimental design and virus purification. C.H. experimental design and virus growth. GvZ experimental design, supervision. S.M. subtomogram averaging. A.S. tomographic reconstruction, particle picking. M.D.M. experimental design, supervision. G.H. data analysis, supervision, paper writing. J.K.L. experimental design, supervision, paper writing. M.B. experimental design, supervision, paper writing.

## Acknowledgments

The cryo electron tomography data was collected at the EMBL Heidelberg Cryo Electron Microscopy Service Platform. We thank EMBL (BT, WH, SM, AS, MB), the Max Planck Society (BT, MS, SW, FECB, SvB, MG, SM, RC, GH, MB), the Max Planck Computing and Data Facility for providing computational resources. We also acknowledge a generous SuperMUC-NG computing allocation at the Leibniz Supercomputing Centre. (MS,SvB,MG,FECB,RC,GH), the Human Frontier Science Program (RGP0026/2017; SvB, GH), the German Ministry of Health (CS), the German Center for Infection Research (CH,MDM) and the Loewe centre DRUID from the Justus Liebig university Giessen (JKL) for funding. BT acknowledges William Wan (Vanderbilt University) for helpful discussions. JKL acknowledges excellent support by Regina Eberle (PEI). RC acknowledges the support of the Frankfurt Institute for Advanced Studies. MS acknowledges support from the Austrian Science Fund *FWF* (*Schroedinger* Fellowship, J4332-B28). The authors are indebted to G. Dobler and R. Wölfel, Bundeswehr Institute for Microbiology, for providing SARS-CoV-2 strain MUC-IMB1. The authors declare no competing interests.

## Materials and Methods

see Supplementary Material - separate document

