## Supplementary for "In situ structural analysis of SARS-CoV-2 spike reveals flexibility mediated by three hinges"

### Supplemental figures and materials

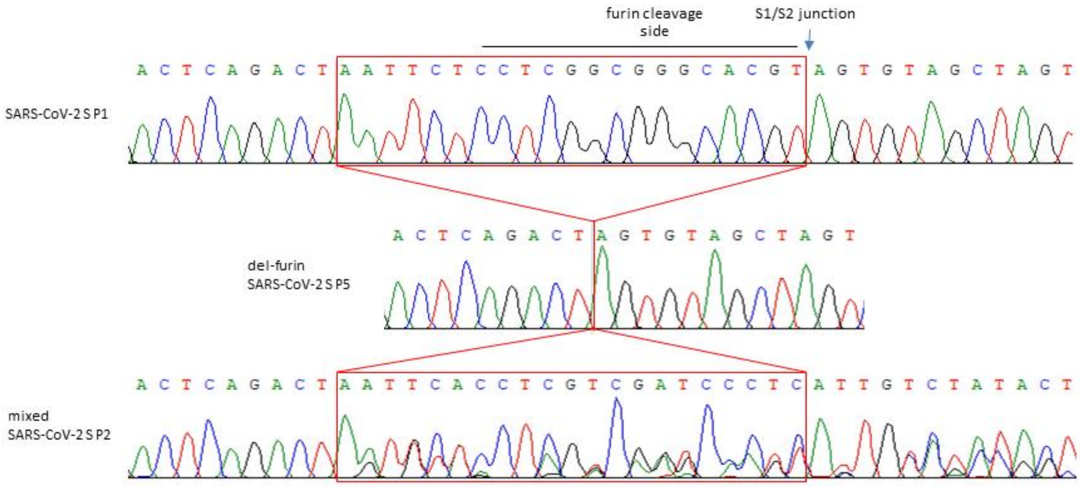


**Figure S1 - Deletion of the furin cleavage site within the viral mRNA in the SARS-CoV-2 S gene.** Sanger sequencing of the SARS-CoV-2 (MUC-1) at passage 1 (P1), 2 (P2) and 5 (P5) propagated on Vero E6 cells as indicated. SARS-CoV-2 S P1 shows the nucleotide sequence spanning the S1/S2 junction region with furin cleavage site of the S glycoprotein. The chromatogram of SARS-CoV-2 passage 5 (P5) shows a 21bp deletion of the furin cleavage side (del-furin). The chromatogram of passage 2 SARS-CoV-2 (P2) shows sequences with and without the 21bp deletion (mixed). Sanger sequencing data from passage 3-4 show exclusively the 21bp deletion (data not shown).

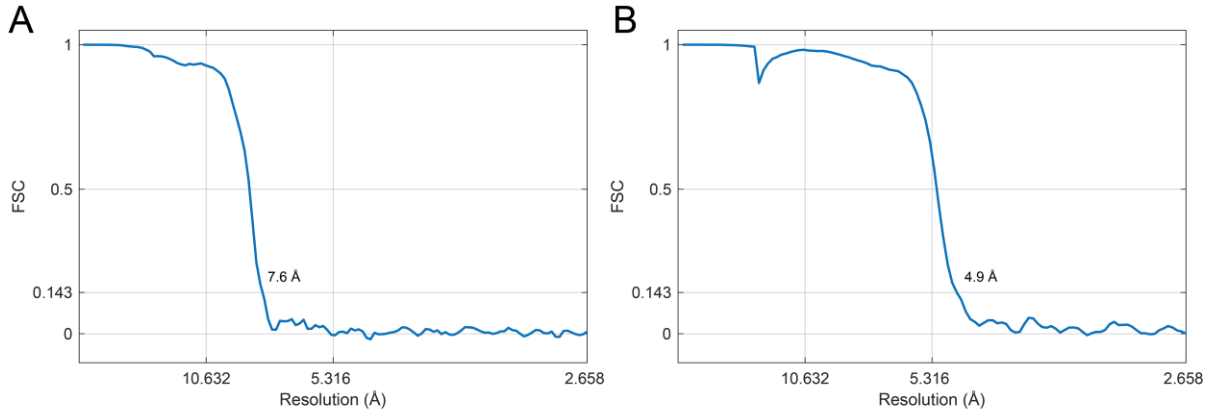


**Figure S2 - Resolution measurement of subtomogram averages.** Fourier Shell Correlation (FSC) curves are shown for the subtomogram averages of the globular domain of spike protein **(A)** and its asymmetric unit **(B)**.

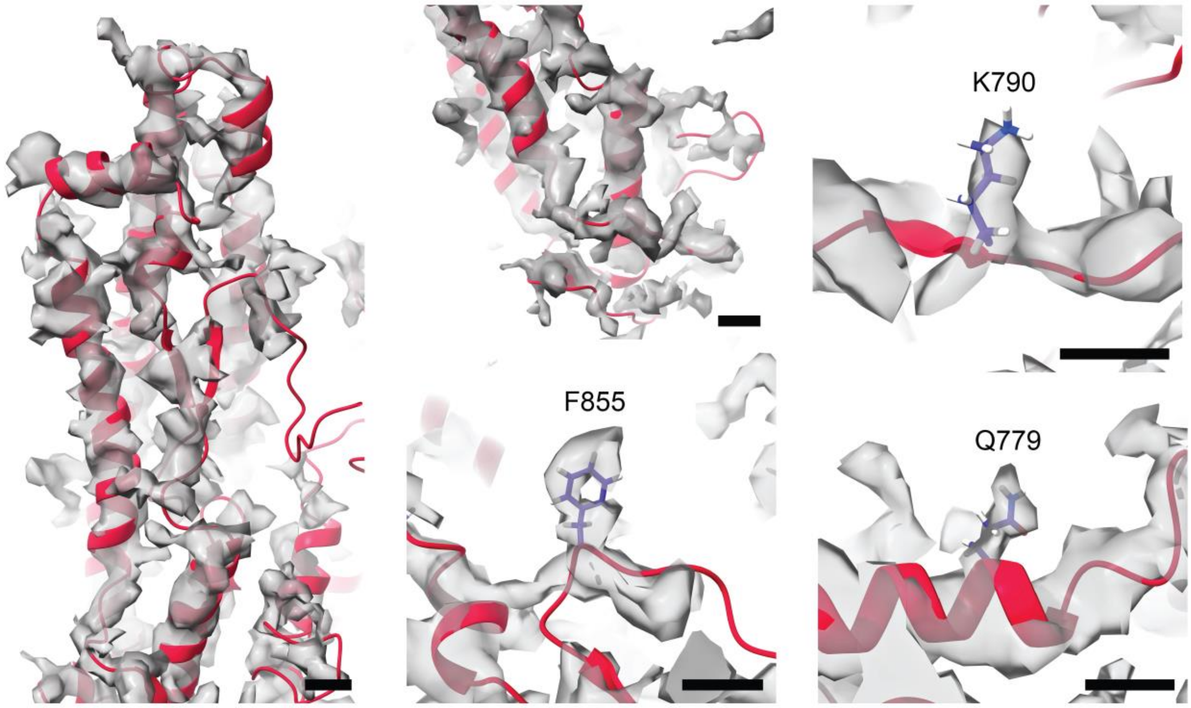


**Figure S3 - Details of the cryoEM map of the asymmetric unit of S.** Averaged electron optical density is shown as isosurface rendered in transparent grey; the MD model was flexibly fitted and is shown as red ribbons. Larger side chains are shown as ball and stick model. Scale bars are 5 Å.

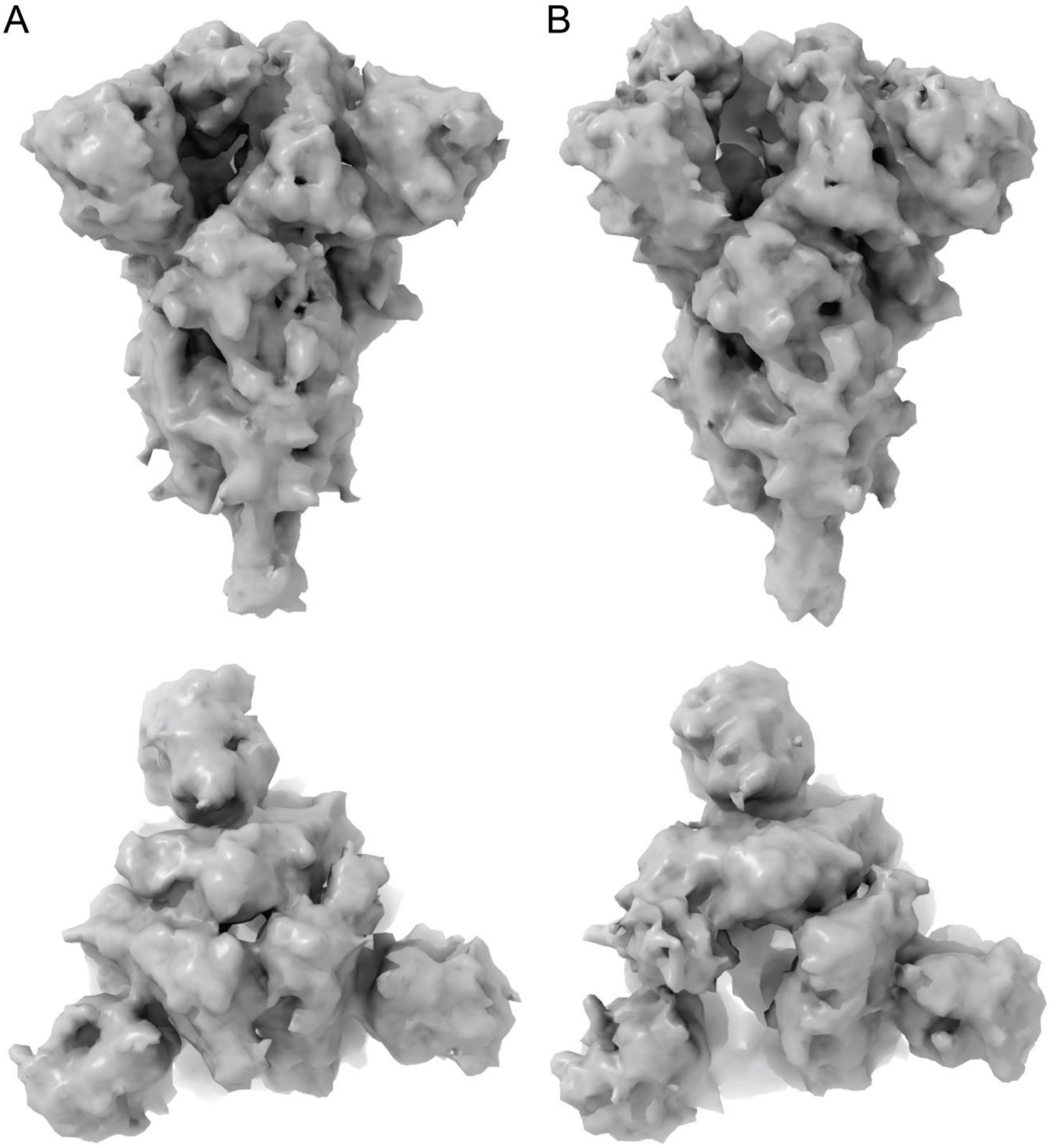


**Figure S4 – Closed and open state conformers. (A)** Side and top view of the subtomogram averaged structure of the fully closed conformation of the S trimer. The depicted structure was obtained from 2x binned data and has a resolution of 9.7 Å. **(B)** Side and top view of the subtomogram averaged structure of S trimer with 1 RBD open. The depicted structure was obtained from 2x binned data and has resolution of 9.7 Å.

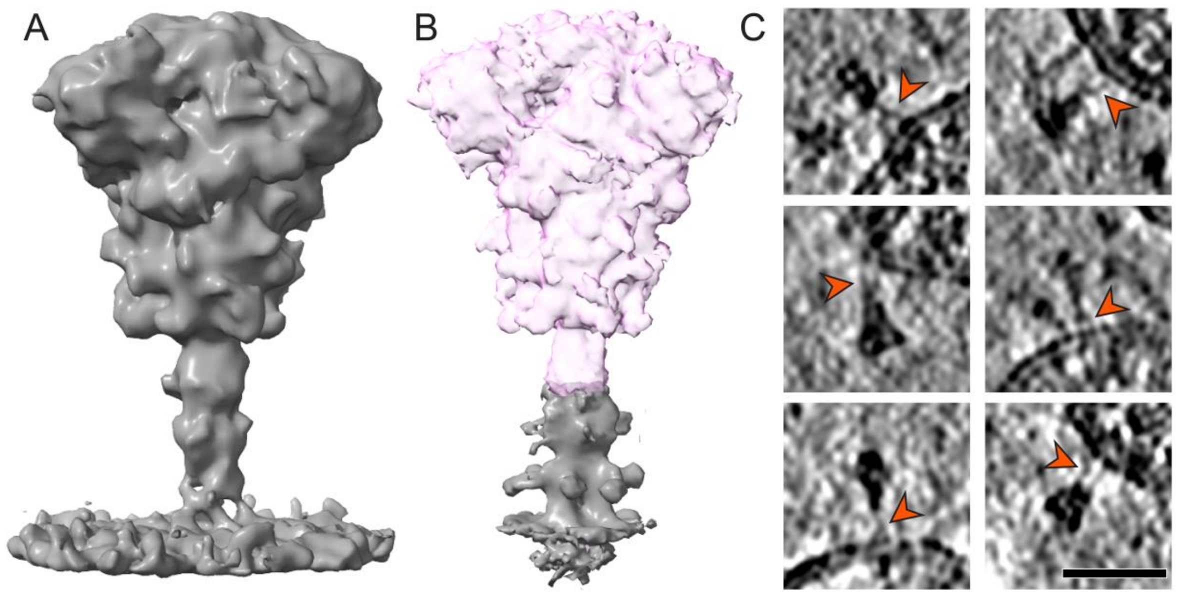


**Figure S5 - Refinement of the lower leg. (A)** Subtomogram averaged structure with 16 Å resolution of the whole body obtained on 2x binned data. **(B)** Structure of the lower leg resolved to 11 Å using focused masking (grey) and superimposed by a separate average of S (pink). **(C)** Examples of unfolded coiled-coils in tomographic slices. Red arrowheads indicated "forking" at hinge positions. Scale bar 30 nm.

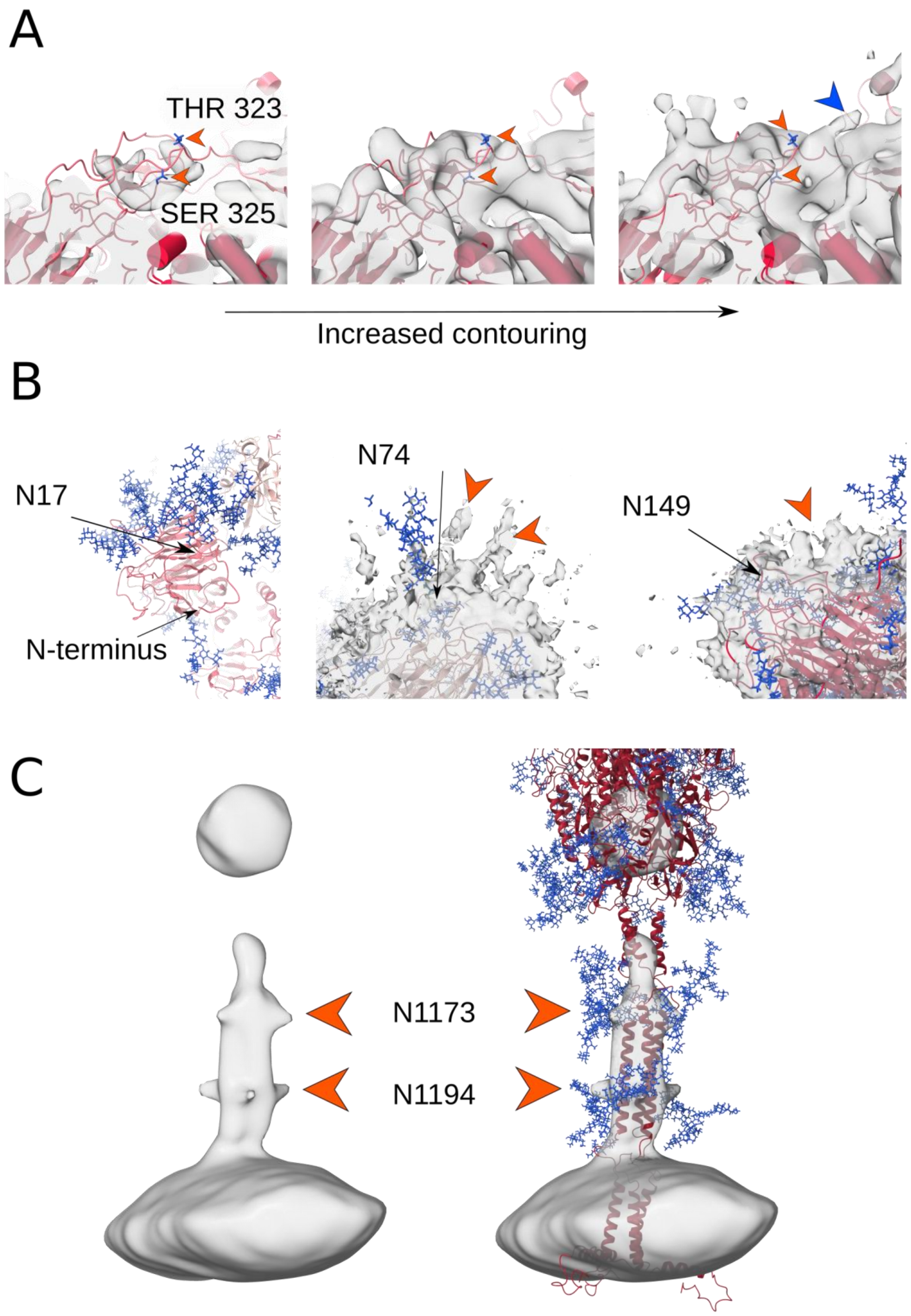


**Figure S6. O- and N-glycosylation of S. (A)** Two potential O-glycosylation sites are marked with orange arrowheads. Maps shown at different contour levels. In comparison to the N-glycosylation sites, these sites lack significant extra density. At the lowest level of contouring an elongated feature becomes visible (blue arrowhead). **(B)** N-glycosylation of the N-terminal domain. The positions of sequons N17, N74 and N149 are indicated with arrows. N17 is located on a flexible N-terminus, likely resulting in very diffuse electron optical density. Unassigned density that could correspond to a glycan is marked with orange arrowheads. **(C)** Average density calculated for the MD trajectory aligned on the lower leg and filtered to the experimental resolution. Left: Due to the flexibility at the hinges, the S head becomes invisible in the averaged density. The lowermost glycosylation sites can be seen protruding from the lower leg. Right: Atomistic model fitted to the density.

**Video S1 - Cryo electron tomography of SarS-CoV-2.** A representative cropped tomogram is shown as constitutive slices along the z-axis. For software used to deconvolve tomograms see [1].

#### Materials and Methods

##### SARS-CoV-2 propagation

SARS-CoV-2 (isolate MUC-1, kind gift of G. Dobler, Bundeswehr Institute for Microbiology) was used for cell infections. It was propagated on Vero E6 cells (ATCC® CRL-1586) with an adsorption of 1 h at 37°C and shaking every 10-15 min. Infected cells were controlled for cytopathic effect every 24h and culture supernatant was usually collected 48h-72h post infection, clarified by centrifugation and was stored in aliquots at -80°C. Virus titer was titrated via TCID50 method according to Kaerber and Spaerman [2].

##### Virus preparation for cryoEM

SARS-CoV-2 identity was verified by Sanger sequencing of a SARS-CoV S gene fragment spanning the furin cleavage site. The RNA genomes of SARS-CoV-2 were isolated from infected Vero cells supernatant using TRIzol LS Reagent (Ambion, Thermo Fisher Scientific, Dreieich Germany) and Direct-zol RNA MiniPrep kit (Zymo research, Freiburg im Breisgau, Germany) according to manufacturers´ instructions and resuspended in 50 µL RNase-free water. Viral cDNA was reverse transcribed using Superscript II RT kit (Invitrogen) with 2 µL vRNA suspension as template and random hexamer primers, according to manufacturer´s instructions. For specific amplification F9newF and F9newR (5´- TAAGGTTGGTGGTAATTATAATTACCTG-3´ and 5´-AAAATAGTTGGCATCATAAAGTAATGGG-3´) primer were used and the respective genomic regions of SARS-CoV-2 S was amplified by PCR and cDNA as template [3]. PCR products were directly sequenced using the following Primers: Wu_24_L and Wu_24_R (5´-TTGAACTTCTACATGCACCAGC-3´ and 5´-CCAGAAGTGATTGTACCCGC-3´) (Eurofins Genomics, Ebersberg, Germany). The furin cleavage site was sequenced at each virus passage. The virus used in this study, corresponding to passage 5 in Vero E6 cells, displayed deletion of residue 2035 to 2055 resulting in the inactivation of the furin cleavage site [4]. For virus purification Vero cells were infected at a MOI of 0.1, the supernatant harvested at 6 hours post-infection and fixed overnight by addition of a final concentration of 4% paraformaldehyde, EM grade (EM sciences). The supernatant was clarified by two times 30 minutes of low-speed centrifugation. The clarified supernatant was placed on top of a 20% sucrose cushion in PBS (w/v) and centrifuged for 90 minutes at 30.000 rpm in a SW41 rotor. The pellet was resuspended in ~100 µl PBS and processed immediately for cryoEM.

##### Western Blot Analysis

Cells were lysed and immunoblotted as previously described [5]. Rabbit anti-SARS-CoV S antibody (1:3,000; ab252690; Abcam) and a rabbit anti-SARS-CoV N antibody (1:1,000; ABIN2214569, Antibody-online) were used for detection of SARS-CoV-2 S and -N respectively. A donkey anti-rabbit IgG-HRP (H&L) polyclonal antibody (1:10,000; Rockland, Gilbertsville, PA) served as secondary antibodies. Peroxidase activity was visualized with an enhanced chemiluminescence detection kit (Thermo Scientific, Bremen, Germany) on ChemiDoc MP Imaging System (Biorad, Dreieich, Germany).

**Cryo electron tomography sample preparation and data acquisition**

Purified virus suspension was mixed with 10 nm protein A-coated gold fiducials (UMC Utrecht). Per sample, 3 µl of the mix was applied onto a glow-discharged 200 mesh copper grid with an R2/2 carbon foil (Quantifoil). Samples were vitrified in liquid ethane using a Vitrobot MarkIV (Thermo Scientific) at 100% humidity and 6°C.

Data was collected from three grids in three microscope sessions at EMBL Heidelberg using a Thermo Scientific Titan Krios G3i electron microscope with a Gatan Bioquantum energy filter and K3 detector. Tilt-series were collected with a dose-symmetric tilt scheme [6,7] using SerialEM software [8]. Tilt controller tilt-range was +/-60 degrees with 3 degree steps. Each tilt image acquisition was preceded by a wait time of 3 seconds, followed by tracking and autofocus. After each tilt-series acquisition, the energy filter zero loss peak was refined, and a new dark reference was acquired. Target focus was changed per tilt-series with a 0.25 µm step over a range from -1.5 µm to -4.5 µm. Tilt images were acquired in superresolution mode with a calibrated physical pixel size of 1.329 Å and a total dose of 3.5 $e^-/Å^2$ over ten frames.

**Image Processing**

**Tilt-series preprocessing and tomogram reconstruction**

The data processing followed the protocol from [7] unless stated otherwise. CTFFind4 [9] was used for CTF estimation and the tilt-series were corrected for dose-exposure [10] using Matlab scripts adapted for tomographic data [11]. The tilt-series were aligned using fiducial-based alignment in eTomo [12]. Gold beads were manually selected and automatically tracked. The fiducial model was corrected in all cases where the automatic tracking failed. The final alignment was computed without solving for any distortions. Two sets of tomograms were reconstructed. The first set consisted of 8x binned tomograms reconstructed using SIRT-like filter option in eTomo. These tomograms were used only for manual selection of virions' centers. A second set of tomograms was reconstructed with 3D-CTF correction using novaCTF [13] with phase-flip correction, astigmatism correction and 15nm slab. Tomograms were binned 2x, 4x and 8x using Fourier3D [14]. The second set was used for subtomogram averaging (SA). SA on binned data was performed with novaSTA [15] while the final refinement on unbinned data was done with STOPGAP [16].

**Subtomogram averaging of the spike**

In total 1094 virions centers were manually picked and their radii estimated. A spherical shape was used to generate initial positions and orientations on the lattice [17]. The sampling distance among the positions on the sphere was 4 nm. Ten iterations of alignment on 8x binned tomograms were run to allow the initial positions to lock on the membrane bilayer. The misaligned positions were removed using the geometry-based procedure described in [7]. The remaining positions were used for ellipsoid fitting, which gave the parameters defining the algebraic equation for conics as well as more precise virion centers and radii. The mean radius of virions obtained in this way was ~40 nm. Five tomograms with lower defocus were used to manually pick the positions of 402 spikes. The initial orientations were generated in the same way as for the positions on the membrane. Ten iterations of alignment on 8x binned tomograms were run to obtain an initial reference of the spike. A new set of starting positions and orientations was generated for each virion by oversampling a sphere using the experimentally determined center and radius which was extended by 15 nm, i.e. the new positions were created roughly in the center of spikes rather than on the membrane. The sampling distance was 4 nm (which roughly corresponds to oversampling of 5-fold). The subtomograms that were too close to a carbon edge were automatically removed. Ten iterations of alignment were run on 8x binned tomograms using the initial reference of the spike. The dataset was subsequently cleaned to remove the subtomograms that shifted to the same positions (distance cleaning), to carbon edge, to the edge of tomograms and/or were overlapping with neighbouring virions. Lastly, subtomograms with low cross-correlation (CC) values were removed from the dataset. As the CC values differ with respect to the tomogram defocus, the CC-based cleaning was done per tomogram. The CC values followed a bimodal distribution and thus the Otsu threshold [18] was used to find the optimal threshold for cleaning. To assure the best possible alignment, i.e. not driven by false-positive particles or the carbon edge, ten iterations of alignment were run on 8x binned tomograms but this time without any starting reference. The positions of the remaining subtomograms were kept but their orientations were set to the initial ones. The subtomograms were split in two halves and processed independently. Four iterations of alignment we run on 4x binned tomograms followed by 3 iterations of alignment on 2x binned data. Prior to the alignment of unbinned data, subtomograms with outlying density values were removed by fitting a normal distribution into the mean density values of subtomograms and discarding those with mean value less or greater than the standard deviation. Five iterations of alignment were run on unbinned data followed by CC cleaning. The final number of particles was 20839. Five iterations of alignment were run, resulting in 7.6 Å structure of the spike. No symmetry was imposed during the alignment.

**Subtomogram averaging of the asymmetric unit**

The positions and orientations from the alignment on 4x binned data were used as a starting point for the asymmetric unit alignment. The spike was split into its asymmetric units which were superimposed, effectively triplicating the number of subtomograms. Four iterations of alignment were run on 2x binned subtomograms. Prior the alignment of unbinned data subtomograms with outlying density values were discarded. Fifteen iterations of alignment on unbinned data were run, followed by CC cleaning and final 5 iterations of alignment. The final average was computed from 65780 particles yielding a 4.9 Å structure.

**Subtomogram averaging of the lower leg**

The average obtained after the alignment on 4x binned data was used to place the mask at the lower leg region and the corresponding positions and orientations were used as starting point for the alignment. In the first iteration on 4x binned data an angular range up to 90 degrees was chosen to allow locking on the lower leg followed by 2 iterations of 30 degrees angular range. The center of rotation was in the knee region of the leg. Three iterations on 2x binned data were run using both novaSTA and STOPGAP with the center of rotation being the lower leg. Simulated annealing in combination with stochastic hill climbing from STOPGAP was used to obtain the structure of the lower leg (see Figure S5B). The structure was resolved to a resolution of 11 Å using 8885 particles and imposing C3 symmetry.

**Distance-based classification**

The orientations and positions found during the alignment on 8x binned data were used to measure the distance of the spike to the membrane. For each spike an intersection of its normal (with inverted direction) and the experimentally determined ellipsoid describing its corresponding virion was found. The distance was measured as the length between the spike center and the intersection with the membrane. Since the lengths of the upper and lower leg are similar this method gives sufficiently precise results also in cases where the spike is bend. However, in some cases the intersection could not be determined. This occurred mostly due to particle misalignment and in rare cases due to strong bending of a spike at one of its hinges. The particles without the intersection point were removed from further classification. Particles with distance less than 9 nm were also discarded as they consisted mostly of misaligned particles that shifted to the membrane during the alignment. The mean distance value was ~19 nm with standard deviation of ~4 nm. Four classes were created based on this: class 1 with distances 9-13 nm, class 2 (13-17 nm), class 3 (17-21 nm), and class 4 with distances larger than 21 nm. The class occupancy was 1560, 4366, 7273, and 5252 particles, respectively. Four iterations of alignment on 8x binned data were run to obtain averages for each class (see Figure 2F). The spike orientations towards the membrane were computed as an angular deviation between the spike normal and an expected membrane normal at the intersection point. The particles with a consistent distance from the membrane (up to the deviation of 2 nm from the mean) and with angular deviation of less than 35 degrees from the membrane normal were chosen for alignment of the whole spike including the leg. The final structure was resolved from 2x binned data to 15 Å (see Figure S5A) using 3193 particles and without imposing any symmetry.

**Classification of closed and open conformers**

Cross-correlation of three possible in-plane orientations of the averaged structure against the previous single particle structures (EMD-21452, EMD-21457) with either the closed or open RBD(s) was computed for each subtomogram to identify the particles with open RBD(s). For partially opened particles the position of the open RBD(s) was subsequently detected de-novo to exclude possible reference bias. Only particles that could be reliably assigned to one of the classes were used for further processing.

**S protein distance distribution and cluster analysis**

We analyzed the organization of S proteins on the viral surfaces in terms of distance and cluster-size distributions. For this analysis, we used the (x,y,z) positions of S proteins on 418 virions. To avoid possible bias in top views, we included S proteins with a polar angle between $\pi/3$ and $2\pi/3$ with respect to the z axis, giving us ~8150 spike positions. We corrected for the geometric restriction in this selection, for variations in virion size, and for the curved shape of the viral surface by repeatedly drawing an equal number of points randomly distributed within a ring around the equator of spheres matching in size. For reference, we also sampled the distribution of equal numbers of hard disks 10 nm in diameter on these spherical waists. For actual, random, and hard-disk S-protein positions, we calculated the distribution of pair distances $r$. The probability ratio for actual and random positions defines a pair-distance distribution function $g(r)$, where a value of one indicates a random distribution of the S proteins on the viral surface. For actual and hard-disk S-protein positions, we also performed a clustering analysis, assigning two proteins (or disks) to the same cluster if their centers are within 20 nm. By comparing the cluster-size distributions for S protein on virions and for hard disks, we assessed the tendency of S proteins to cluster beyond what is expected to occur at random.

**Molecular dynamics simulations**

We embedded four full length S proteins in a membrane of ER-like composition. After adding aqueous solvent, the simulation system contained ~4.1 million atoms. An all-atom model was constructed based on a single particle cryoEM structure (PDB ID: 6VSB, [19]), supplemented with a coiled-coil model of the stalk and transmembrane domain (TMD) and full glycosylation pattern [20](Figure 3A). We note that the simulation model described here had been generated independently from the tomographic data and before they were available. Although the simulation model and the EM maps are overall remarkably consistent, in particular concerning the positions of the hinges, there are some minor deviations. Notably, the short coiled coil forming the upper leg was modeled as left handed. Even though this does not affect the conclusions here, it will be interesting to explore the effect on stability of changing the handedness. We note further that the relatively high effective density of S proteins in the simulation system at ~15 nm spacing sterically restricted the mobility, thus preventing large bending angles.

MD simulations of 2.8 microseconds at constant pressure (1 bar) and temperature (310 K) were performed with particle-mesh Ewald electrostatics, a time step of 4 fs and nonbonded interactions cut off at 1 nm using GROMACS 2019.6[21]. Structures were saved at intervals of 10 nanoseconds and used to identify the flexible hinges in the structure and to calculate density maps for as a basis for quantitative comparisons with cryoEM.

**Calculation of the bending angles**

Vectors defining hinge angles were calculated as follows: Hip: Head center of mass - hip center of mass - knee center of mass; knee: hip center of mass - knee center of mass - ankle center of mass; ankle: knee center of mass, ankle center of mass - TMD center of mass. Angle probability densities were estimated using kernel estimation as implemented in SciPy [22].

**Generation of density maps**

Trajectories of all four S proteins were aligned on the head domain. An electron density map was subsequently calculated using Gromaps [23] with the same grid spacing as used in the experiment (0.1329 nm) and spread parameter sigma adjusted to reproduce the experimental resolution.

**Density-guided simulations**

As an initial step maps were superimposed onto one of the S proteins in a simulation box using ChimeraX [24]. Density guided simulations were performed using a dedicated tool in GROMACS 2020.2 [25]. Atoms belonging to S (excluding the TMD domain and including all glycosylation sites) were used as a fitting group. The density cross-correlation was used to quantify agreement between the experimental and calculated density maps, with a scaling factor of 2 x $10^{10}$ kJ/mol. Forces acting on atoms were applied every 100 steps and adaptive scaling was used.

**Rigid body fitting to raw tomogram data**

S proteins were segmented manually using ImageJ [26]. Snaphots of S proteins taken from the MD simulation were selected to best match the geometry of the hinges, centered in the density and rigid-body fitted using ChimeraX [24].

**Rigid body fitting to subtomogram averaged maps**

Rigid body fitting of the single particle model (PDB ID 6VXX) was performed in ChimeraX [24] using Fit in Map tool.

**Visualization**

All subtomogram averaged maps shown were post-processed in RELION using automatic B-factor sharpening [27]. The raw tomograms visualized in the Figures were deconvolved using tom_deconv [1].

14. https://github.com/turonova/Fourier3D

15. https://github.com/turonova/novaSTA

16. https://github.com/williamnwan/STOPGAP

17. Förster, F., Medalia, O., Zauberman, N., Baumeister, W. & Fass, D. Retrovirus envelope protein complex structure in situ studied by cryo-electron tomography. *P. Natl. Acad. Sci. USA* **102,** 4729-4734 (2005).

18. Otsu, N. A threshold selection method from gray-level histograms. *IEEE Trans. Sys. Man. Cyber.* **9** (1), 62-66 (1979).

19. Wrapp D. et al. Cryo-EM structure of the 2019-nCoV spike in the prefusion conformation. Science 367, 1260-1263 (2020)

20. Watanabe Y., Allen J. D. Wrapp D., McLellan J. S., Crispin M. Site-specific glycan analysis of the SARS-CoV-2 spike. Science eabb9983 DOI: 10.1126/science.abb9983
